# Epigenomics identifies three sources of DNA methylation in *Streptococcus mutans* UA159

**DOI:** 10.64898/2026.04.07.717064

**Authors:** Matthew Barbisan, Daniel Kim, Sadie Gavriela Drucker, Michelle Lee, Jonathon L. Baker

**Author notes:** Corresponding Author: JLB.

## Abstract

DNA methylation is a widespread but incompletely characterized regulatory feature of bacterial genomes. While restriction–modification systems represent well-studied sources of DNA methylation, the full complement of methyltransferases shaping bacterial epigenomes and their physiological consequences remain poorly understood. Here, we used Oxford Nanopore sequencing to comprehensively map DNA methylation in the model oral pathogen *Streptococcus mutans* UA159. Genome-wide analysis identified extensive N6-methyladenosine (6mA) modification and revealed three predominant methylation motifs. Using targeted deletion mutants, we demonstrate that methylation at GATC sites is mediated by the conserved DpnII restriction–modification system, while a novel bipartite CGANNNNNNNTCY/RGANNNNNNNTCA motif is methylated by the HsdM component of the type I Hsd restriction–modification system. The remaining 6mA sites corresponded to a CTGNAG/CTNCAG motif, defining the activity of a third methyltransferase. Genetic and epigenomic analyses identified SMU.43 as the enzyme responsible for this modification, which we designate DnmA, a novel orphan adenine methyltransferase with homology to regulatory methyltransferases rather than defense-associated systems. Functional characterization of single and double mutants revealed that distinct methylation systems differentially influence biofilm formation and antagonistic interactions with the commensal, *Streptococcus sanguinis*. Notably, loss of *dnmA* reversed biofilm and aggregation defects associated with deletion of *dpnII*, indicating epistatic interactions between methylation pathways. Together, this study resolves the major sources of DNA methylation in *S. mutans* UA159, identifies a novel regulatory methyltransferase, and highlights the utility of nanopore sequencing for bacterial epigenome discovery. These findings expand our understanding of bacterial DNA methylation and suggest that epigenomic enzymes may represent targets for modulation of microbial physiology and virulence.

## Main Text

DNA/RNA modifications (i.e., noncanonical bases; e.g., methylation, inosine, pseudouridine, etc.) play major roles in the physiology of all kingdoms of life and viruses. Compared to their roles in eukaryotes, these modifications and associated processes (collectively termed the epigenome for DNA and the epitranscriptome for RNA), and their effects, are less broadly understood and appreciated in bacteria, despite doubtlessly having major impacts on microbial communities, and consequently, human health (Oliveira 2021; Höfer and Jäschke 2018). DNA modifications in bacteria were initially identified in the context of restriction-modification defense systems to distinguish self from non-self (Luria and Human 1952). These systems are widely distributed and have a profound impact, allowing bacteria to defend against phages. Contemporary work examining bacterial epigenomics is revolutionizing our understanding of the extent and physiological relevance of bacterial DNA modifications in other contexts. Bacteria exhibit at least 3 types of DNA methylation: N6-methyladenosine (6mA), 5-methylcytosine (5mC), and N4-methylcytosine (4mC)(Figure 1A), playing major roles in physiology, antibiotic resistance, and virulence (Oliveira 2021). For example, it was recently discovered that methylation plays roles in *Salmonella* motility and oxidative stress response (Zhang et al. 2023; Bourgeois et al. 2022), phase variation in *Streptococcus pneumoniae* (Kwun et al. 2023), and gene expression and autoaggregation in *Streptococcus mutans* (Zhao, Dufour, Zhong, et al. 2025; Zhao, Dufour, Ghobaei, et al. 2025). Traditional study of modified and noncanonical bases involved inspection of suspected sites on an individual site basis with biochemical assays or specialized sequencing methods to detect a single modification (e.g., bisulfate sequencing). Conversely, emerging third-generation sequencing technologies (TGS; e.g., Oxford Nanopore Technologies [ONT] and PacBio) are poised to revolutionize this field. Second-generation sequencing technologies, such as Illumina, that use sequencing-by-synthesis, do not sequence the original, native nucleic acid molecule. This means that any epigenomic and epitranscriptomic information is lost before or during the sequencing process. However, emerging TGS technologies do sequence native nucleic acid molecules and can, for the first time, detect multiple DNA and RNA modifications *en masse* on a genome, transcriptome, or microbiome-wide scale.

**Figure 1:**
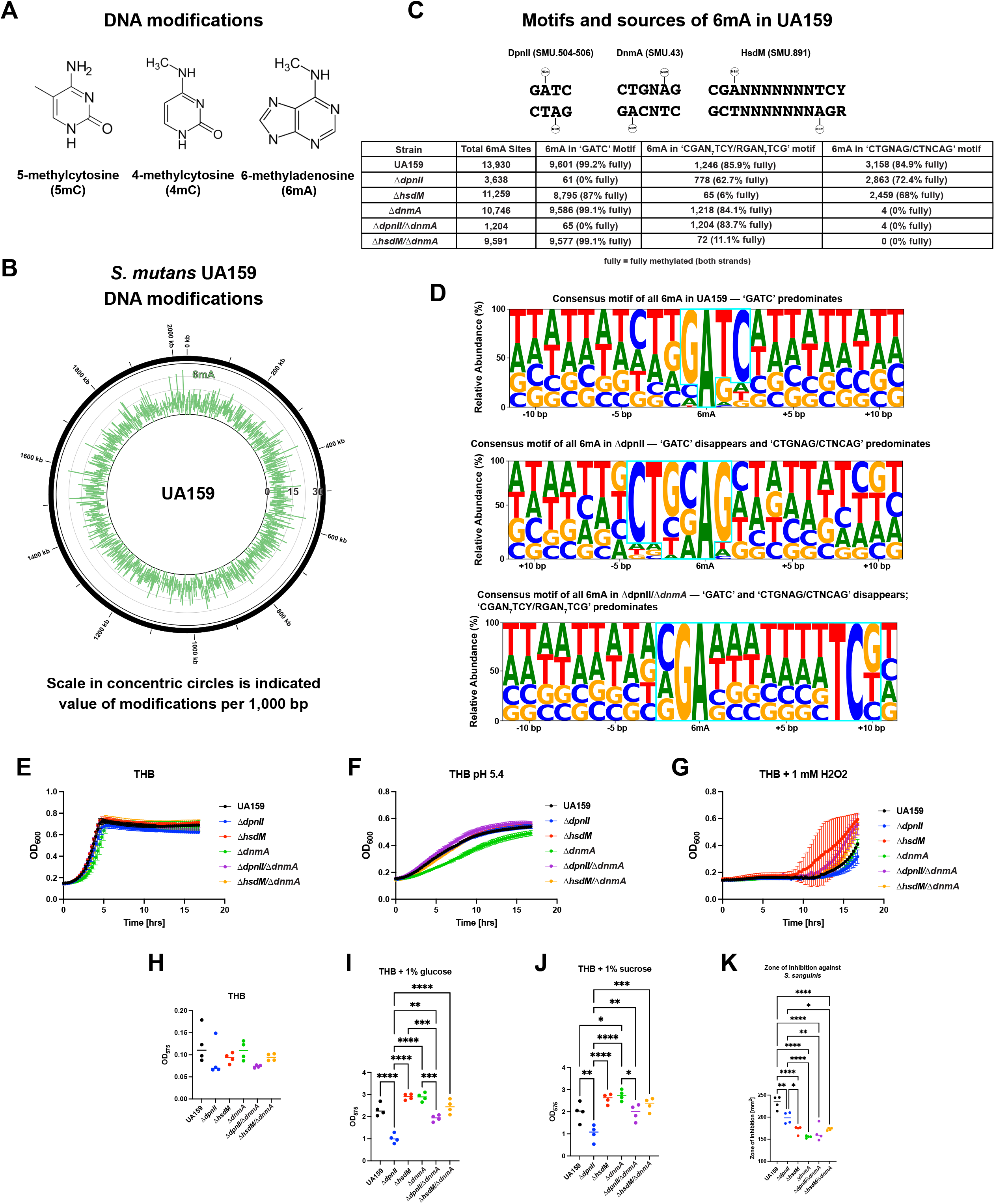
**Panel A:** Structures of the DNA modifications queried for in *S. mutans* UA159 in this study. **Panel B:** Plot of the locations and density of the 6mA detected in the UA159 genome. The scale in the concentric circles is the indicated value of the indicated modification per 1000 bp window. **Panel C:** Structure of the 3 6mA motifs identified in UA159, along with the responsible genes, and a table describing the number of 6mA found in each motif in the indicated strain. Value in parenthesis indicates the percentage of those 6mA in a fully-methylated site (i.e., the motif is methylated on both DNA strands). **Panel D:** Consensus logos illustrating the prevalence of each nucleotide in the indicated window adjacent to all 6mA in the indicated strain. **Panels E-G:** Growth curves of the indicated strains in either THB (Panel E), THB buffered to pH 5.4 (Panel F), or THB + 1 mM H_2_O_2_ (Panel G). **Panels H-J:** Scatter plots illustrating the biofilm formation determined by a crystal violet assay of the indicated strain under the indicated condition. Asterisks indicate statistical significance (**, P < 0.01, ****, P < 0.0001) as determined by a Tukey’s Honestly Significant Differences test following a one-way ANOVA. **Panel K:** Scatter plot illustrating the zone of inhibition of *S. sanguinis* growth from the indicated strain of *S. mutans*. Asterisks indicate statistical significance (*, P < 0.05, ***, P < 0.001) as determined by a Tukey’s Honestly Significant Differences test following a one-way ANOVA.

In this study, the epigenome of the model pathogen *Streptococcus mutans* UA159 was analyzed using ONT sequencing, allowing for simultaneous detection of 6mA, 5mC, and 4mC in the DNA. *S. mutans* is a major etiologic agent of dental caries, the most common chronic infectious disease globally (Pitts et al. 2017). *S. mutans* has also been widely used as a model for Gram-positive bacteria, due to its genetic tractability, which has also led to its utility as an expression system to examine natural products synthesized by anaerobic bacteria (Lemos et al. 2019; Hao et al. 2019). Here, the archetype strain *S. mutans* UA159 was grown to mid-log phase in Todd Hewitt broth (THB). The DNA was then harvested from pelleted cells and sequenced on an Oxford Nanopore MinION. With a minimum coverage of 10 and a minimum percent modified of 50%, UA159 contained 13,930 6mA (Figure 1B, Supplemental Table S1). 9,601 (69%) of the 6mA were within a GATC motif, 1,246 (9%) were within in a CGANNNNNNNTCY motif (or its reverse complement sequence RGANNNNNNNTCG), 3,158 (23%) were within a CTGNAG motif (or its reverse complement CTNCAG)(Figure 1C-D, Supplemental Tables S1 and S2). The counts of the 6mA listed in the previous sentence to add up to 14,005, which is more than the total number of 6mA in the genome (13,930) due to overlap of the motifs in some cases (e.g., C**GATC**NNNNNTCY). No 4mC or 5mC were detected in UA159.

The DpnII Type II restriction-modification (R-M) system (SMU.504, SMU.505, and SMU.506 in the *S. mutans* UA159 genome) is understood to methylate adenines at the N6 position within the sequence GATC (Lacks et al. 2000). The DpnII machinery (SMU.504-506) is highly conserved in *S. mutans* in terms of both carriage of the genes and homogeneity of the protein sequences. It appeared in 222 of the 244 strains included in our previous pangenome analysis (Baker et al. 2022) with the DpnII protein sequences having a combined homogeneity index (Eren et al. 2021) of 0.998%. An *S. mutans* strain lacking the DpnII R-M system was recently characterized and displayed reduced methylation (analyzed in bulk), increased susceptibility to oxidative stress, eDNA-dependent autoaggregation, and decreased biofilm formation (Zhao, Dufour, Zhong, et al. 2025). Loss of *dpnII* also led to differential regulation of 60 genes, in part through methylation in the promoter regions impacting transcription (Zhao, Dufour, Ghobaei, et al. 2025). A similar strain (Δ*dpnII*) was generated for this study (described in Materials & Methods). The Δ*dpnII* strain did not have a significantly different growth phenotype compared to UA159 in THB, THB buffered to pH 5.4, or THB + 1 mM H_2_O_2_ (Figure 1E-G). However, similar to what was observed by Zhao et al., the loss of *dpnII* did result in a significant decrease in biofilm formation in THB supplemented with glucose or sucrose (Figure 1H-J)(Zhao, Dufour, Zhong, et al. 2025). The Δ*dpnII* strain also produced a smaller zone of inhibition in a deferred antagonism assay against *Streptococcus sanguinis*, a health-associated organism that *S. mutans* ecologically competes with (Figure 1K). When subjected to the same DNA sequencing and modification analysis as UA159, the Δ*DpnII* strain contained only 3,638 6mA sites using the same cutoffs of 50% modified bases and a coverage of 10 applied for UA159 (Figure 1C, Supplemental Table S1). Since DNA is double stranded, when a methylated motif is methylated on both strands, it is referred to as fully-methylated, whereas when the motif is only methylated on one strand, it is referred to as hemi-methylated. Crucially, there were zero fully methylated 6mA GATC motifs in Δ*dpnII* identified by modkit, which is in agreement with DpnII methylating adenosines within the GATC motif (Supplemental Tables S1 and S2). In Δ*dpnII*, all 61 remaining 6mAs at GATC motifs (identified by custom scripts, not modkit, see UA159-epigenome repository on GitHub) represented hemi-methylated motifs, whereas in UA159, 99.2% of the GATC sequences were fully methylated (Figure 1C). These hemi-methylated GATC sequences in Δ*dpnII* also all overlapped with CGANNNNNNNTCY/RGANNNNNNNTCA sequences, indicating the methylation is from the other motif and related machinery, not DpnII. Both CGANNNNNNNTCY/RGANNNNNNNTCA and CTGNAG/CTNCAG 6mA motifs were identified by modkit in Δ*dpnII*, similar to what was observed in UA159.

The bipartite CGANNNNNNNTCY/RGANNNNNNNTCA motif identified by modkit in UA159 and Δ*dpnII* was reminiscent of the recognition sequences of the type I *hsd* restriction-modification system. This system has not been previously characterized in *S. mutans*, but has been studied rather extensively in other species, including *S. pneumoniae*, where it is responsible for phase-variation (Li et al. 2016), and *S. pyogenes*, where it is a major factor blocking genetic manipulation (Finn et al. 2021; Bjånes et al. 2024; DebRoy et al. 2021). To determine whether the methylations at CGANNNNNNNTCY/RGANNNNNNNTCG were indeed a product of the *S. mutans hsdI* system, a deletion mutant of *hsdM* (SMU.891) was examined using the same ONT sequencing methods applied above. Indeed, the Δ*hsdI* strain had only 65 6mA within CGANNNNNNNTCY/RGANNNNNNNTCG, with only 6% of those fully-methylated in comparison to UA159, which had 1,246 6mA in this motif with 86% of those fully-methylated (Figure 1C, Supplemental Tables S1 and S2). The remaining hemi-methylated *hsdM* sites also overlapped with CTGNAG/CTNCAG, again indicating that these were associated with the other motif and machinery. Despite not being reported by modkit in the Δ*hsdM* strain, CTGNAG/CTCNAG was still methylated with 2,459 6mA associated with this motif. Like Δ*dpnII*, Δ*hsdM* did not have a different planktonic growth phenotype compared to UA159 in THB, THB pH 5.4, or THB + 1 mM H_2_O_2_, and also exhibited reduced inhibition of *S. sanguinis* (Figure 1E-G,K). Unlike Δ*dpnII*, deletion of Δ*hsdM* did not have a significant impact on biofilm formation (Figure 1H-J). The *S. mutans* UA159 *hsd* system is not contiguous, with *hsdM* and *hsdS* (SMU.892) being separated from *hsdR* (SMU.897) by a putative anticodon nuclease *prrC* (SMU.893) (Bacusmo et al. 2018) and relB/relE toxin-antitoxin system (SMU.895-896) (Tian et al. 2019; Shields and Burne 2016). Interestingly, *hsdRMS* was not widespread across *S. mutans*, as the *hsdM* was only found in 18 strains, *hsdS* in 13 strains, and *hsdR* in 27 strains of the 244 strains in the published *S. mutans* pangenome (Baker et al. 2022). These genes had combined homogeneity indices of 0.994 for *hsdM*, 0.745 for *hsdS*, and 0.993 for *hsdR*. This agrees with the function of *hsdS* as the DNA specificity subunit, and having a much lower identity (i.e., so it can bind to different DNA consensus sequences). Across the *S. mutans* pangenome, there were at least 8 different configurations of the *hsd* genes, with some missing components or having duplicate *hsdS* genes (Supplemental Table S3), reminiscent of the *S. pneumoniae hsd* system where the multiple copies facilitate recombination and therefore phase variation. All but one, however, were still adjacent to *relB/relE*.

Since the majority of the remaining (i.e., not explained by DpnII or HsdM) 6mA sites found in the *S. mutans* strains examined here were associated the CTGNAG/CTNCAG motif, it is likely that this represented an unidentified third source of methylation. SMU.1979 is predicted to be an adenine-specific DNA N6-methylase and is found within the nine gene *comY* operon in all but one *S. mutans* genome in the pangenome with a homogeneity of 99.8%. A deletion mutant of SMU.1979 also reduced competence to 40% of the level observed in UA159 (Merritt, Qi, and Shi 2005). A deletion mutant of this SMU.1979 was examined using nanopore sequencing. However, all 3 motifs described above were still present in ΔSMU.1979, indicating that SMU.1979 is not responsible for the 6mA in CTGNAG/CTNCAG in the UA159 genome under the conditions tested (Supplemental Tables S1 and S2). SMU.43 is also predicted to be an adenine-specific DNA methyltransferase, present in 60 *S. mutans* genomes across the pangenome with >99% homogeneity. A ΔSMU.43 strain was generated and this strain had no 6mA within the CTGNAG/CTNCAG motif, defining the recognition motif of this novel methyltransferase. SMU.43 contains a YeeA motif, similar to the recently described *Bacillus subtilis* DnmA, indicating its activity may function in regulation of gene expression, rather than defense against foreign DNA (Nye et al. 2020). Therefore, we are designating SMU.43 as *dnmA*. Δ*dnmA* had similar growth to UA159 in THB and THB + 1mM H_2_O_2_ and had slightly impaired growth compared to UA159 and the other methylation mutants in THB pH5.4, indicating slightly reduced acid tolerance, a virulence factor of *Streptococcus mutans*. Compared to UA159, Δ*dnmA* was not defective in biofilm formation during growth in either glucose or sucrose. Δ*dnmA* did produce a significantly smaller zone of inhibition against *S. sanguinis* in the deferred antagonism assay.

To examine the impact of removing multiple sources of methylation, double mutants Δ*dpnII/*Δ*dnmA* and Δ*hsdM/*Δ*dnmA* were also constructed. In both cases, the methylation at the two expected cognate motifs was lost. Interestingly, the Δ*dpnII/*Δ*dnmA* double mutant reversed the biofilm-deficient phenotype and coaggregation phenotype observed in the Δ*dpnII* single mutant. Both double mutants had a similar reduction in the zone of inhibition produced against *S. sanguinis* compared to the Δ*dnmA* single mutant. Continued research examining the role of these methylation systems in regulating the *S. mutans* transcriptome and determining how deletion of *dnmA* reverses the phenotypes of *dpnII* is currently in progress. Overall, this study identifies DnmA as a novel streptococcal orphan methyltransferase as well as elucidates the recognition sequence of the UA159 HsdM methyltransferase. Furthermore, this study illustrates the power of nanopore sequencing to analyze bacterial epigenomes and provides resources and instructions to other research groups on how to conduct this type of analysis. Since the types of methylation found in bacteria are different than those found in eukaryotes, the bacterial DNA modification enzymes could represent a therapeutic target.

## Materials and Methods

### Bacterial strains and growth conditions

All *Streptococcus* strains were maintained on Todd-Hewitt broth (THB) agar plates (BD/Difco, Franklin Lakes, NJ) at 37°C in a 5% (vol/vol) CO_2_– 95% air environment. The *S. mutans* strain UA159 has been previously described (Ajdić et al. 2002). Δ*hsdM* and ΔSMU.1979 were generated previously as part of a library of single-gene deletion mutants in *S. mutans* UA159 (Quivey et al. 2015). The Δ*hsdM* ONT sequencing described below confirmed the expected mutation of the *hsdM* open reading frame and no SNPs, indels, or SVs were identified in Δ*hsdM* compared to our UA159 strain. **DpnII deletion strain (**Δ***dpnII*):** A construct was synthesized by Integrated DNA Technologies that included 500 bp upstream of SMU.504, followed by the *ermB* (erythromycin resistance gene) open reading frame, followed by 500 bp downstream of SMU.506. This construct was transformed into UA159 as described in (Lau et al. 2002) and transformants were selected by plating on THB + 5 µg mL^-1^ erythromycin. Correct insertion and orientation of the construct was confirmed by the ONT sequencing described below. The sequencing detected no SNPs, indels, or SVs in the Δ*dpnII* strain. **DnmA deletion strain (**Δ***dnmA*):** A construct was synthesized by Integrated DNA Technologies that included 300 bp upstream of *dnmA* (SMU.43), followed by the kanamycin resistance gene (kanR) open reading frame, followed by the 400 bp downstream of *dnmA*. This construct was transformed into UA159 as described in (Lau et al. 2002) and transformants were selected by plating on THB + 1 mg mL^-1^ kanamycin. Correct insertion and orientation of the construct was confirmed by the sequencing described below. The ONT sequencing of Δ*dnmA* did identify that the strain had a A -> C mutation with 97% frequency at position 1,181,710 in SMU.1243 (low-temperature requirement protein A), causing an in-frame A -> D mutation. Additionally, there was a recombination of the *aroG1* (SMU.1836) and *aroG2* (SMU.1837) genes to form a hybrid protein. Further research on the identified SNP and the *aroG1/G2* recombination event is in progress, but the fact the Δ*hsdM/*Δ*dmnA* and Δ*dpnII/*Δ*dnmA* double mutants did not have these mutations and still had the same loss of 6mA at the CTGNAG/CTNCAG motif demonstrates that the methylation loss in Δ*dnmA* is not due to these second site mutations. **DpnII/DnmA double mutant (**Δ***dpnII/***Δ***dnmA*):** The same construct described in the previous section to generate the Δ*dnmA* strain was transformed in the same manner as described above, except into the Δ*dpnII* strain instead of UA159. Transformants were plating on THB + 1 mg mL^-1^ kanamycin and + 5 µg ml^-1^ erythromycin. Correct insertion and orientation of the constructs was confirmed by the ONT sequencing described below. The sequencing also identified no unexpected SNPs, indels, or SVs compared to our UA159 strain. **HsdM/DnmA double mutant (**Δ***hsdM/***Δ***dnmA*):** The same construct described above to generate the Δ*dnmA* strain was transformed in the same manner as described above, except into the Δ*hsdM* strain instead of UA159. Transformants were plating on THB + 1 mg mL^-1^ kanamycin and + 5 µg mL^-1^ erythromycin. Correct insertion and orientation of the construct was confirmed by the sequencing described below. The sequencing also identified no unexpected SNPs, indels, or SVs compared to our UA159 strain.

### Oxford Nanopore Sequencing of genomic DNA

For gDNA sequencing, colonies of UA159, Δ*dpnII*, Δ*hsdM*, Δ*dnmA*, Δ*dpnII/*Δ*dnmA*, and Δ*hsdM/dnmA* were inoculated in 5 mL liquid THB media and cultured overnight) at 37°C in a 5% (vol/vol) CO_2_–95% air environment. The following day, the inoculums were centrifuged at 18,000 x g for 10 min to form cell pellets and the supernatant media was gently aspirated. Bacterial chromosomal DNA was isolated from the cell pellets using the Zymo Quick-DNA Fungal/Bacterial Miniprep Kit according to the manufacturer’s instructions. Sequencing libraries for the strains were prepared using the Ligation sequencing gDNA – Native Barcoding Kit 24 V24 (Oxford Nanopore Technologies, Inc.), according to the manufacturer’s instructions. The barcoded DNA libraries for each strain was sequenced with a MinION Flow Cell on a MinION Mk1B sequencer (Oxford Nanopore Technologies, Inc.), with sequencing runs monitored using MinKNOW v25.03.9 (Oxford Nanopore Technologies, Inc.). The resulting POD5 files which were used for downstream analysis.

### Variant analysis

To confirm the expected sequence of each strain and identify any SNPs, indels, or structural variants, FASTQ read files were extracted from the POD5 files and imported into CLC Genomics Workbench v26.0.1. The read files were mapped against our UA159 reference sequence using the “Map Long Reads to Reference” function. The resulting mapping file was then analyzed with the “Basic Variant Detection” and “InDels and Structural Variants” tools and the results were manually inspected.

### Basecalling and identifying modified bases

While MinKNOW provides the ability to basecall the POD5 files directly during data acquisition, using the super-high accuracy basecalling model in addition to the modified bases parameters results in large processing times. Because of this, the POD5 files were uploaded to the OHSU ARC high-performance compute cluster where Dorado v1.0.2 (Oxford Nanopore Technologies basecalling software) was used to process the raw POD5 reads. The Dorado basecalling model dna_r10.4.1_e8.2_400bps_sup@5.0.0 was used for processing and filtering the DNA POD5 reads with a q-score ≥ 9. The modified bases (methylation sites for DNA) were identified using the modified base models dna_r10.4.1_e8.2_400bps_sup@v5.0.0_ 6mA@v1 and …_4mC_5mC@v1 for the 6-methyladenine (m6A), and 4- and 5-methylcytosine (m4C and m5C) sites, respectively. The resulting BAM files contained the annotations for modified base predictions and the associated probabilities, which was further analyzed by modkit (v0.5.0) (https://github.com/nanoporetech/modkit). The BAM files generated by modkit were used as input for a custom pipeline (https://github.com/jonbakerlab/UA159-epigenome) inspired by open-source ont-methylation software (Galeone et al. 2025). This pipeline mapped the BAM file to a supplied FASTA file of the UA159 genome and produced a bedMethyl file from the pileup function in modkit and then extracted and filtered the data into a list of methylated positions found in each strain.

### Growth Curves

Overnight cultures of the indicated strains were normalized to the OD_600_ of the lowest density culture in the comparison using the indicated growth media. Growth curves were performed in either 96-well or 384-well clear plates (Corning, Inc.). For growth curves in 96-well plates, 10 µl of the overnight cultures were added to 200 μl of the indicated growth media with or without indicated compounds. For growth curves using 384-well plates, 2.5 µl of the overnight cultures were added to 50 µl of the indicated growth media with or without the indicated compounds. Plates were sealed with Breathe-Easy sealing membrane (Diversified Biotech) and growth curves were performed without the plastic plate lid. Growth was monitored using an Infinite M Plex (Tecan Group, Ltd.).

### Deferred antagonism (i.e., “competition”) assay

The deferred antagonism assay was performed as previously described (Uranga et al. 2021). Briefly, 8 mL of overnight cultures of the initial colonizer was spotted onto THB 1% agar and incubated overnight at 37°C under 5% CO2/95% air. The following day, the plates were sterilized using the sterilization setting (90 s) in a GS Gene Linker UV Chamber (Bio-Rad, Inc.). 500 µL of overnight cultures of *S. sanguinis* SK36, were added to 5 mL molten THB 0.75% agar that had been cooled to 40°C, and this was used to overlay the plates with the *S. mutans* colonies. The agar overlay was allowed to solidify at room temperature, and then the plates were incubated overnight at 37°C under 5%CO2/95% air. The following day, zones of inhibition were measured using a ruler.

### Biofilm assay

Biofilms of the indicated strains were grown in overnight THB, THB + 1% glucose, or THB + 1% sucrose in 96 well plates at 37°C in a 5% (vol/vol) CO_2_–95% air environment. The following day, the supernatant was aspirated, and the biofilms were washed twice by gently submerging the plate in distilled water. Plates were then dried upside-down on absorbent paper. Adherent biofilms were then stained with 100 µL 0.1 crystal violet solution for 15 minutes. The plates were then washed twice by submersion in distilled water. The bound dye was extracted for 1 hour in 200 µL of 500 mM acetic acid. The eluted dye was then transferred to a new plate, and optical density at 575 nm was measured using a Tecan Infinite M Plex.

## Data Availability

The raw sequencing data (POD5 and FASTQ files) will be made available in the final peer-reviewed publication. All code and more detailed methods associated with this project are available at https://github.com/jonbakerlab/UA159-epigenome.

## Acknowledgements

This research was supported by NIH/NIDCR R00-DE029228 (J.L.B.), R25-DE032536 (S.D., M.L.), and by Oregon Health & Science University. The research reported in this publication also used computational infrastructure supported by the Office of Research Infrastructure Programs, Office of the Director, of the National Institutes of Health under Award Number S10-OD034224. The content is solely the responsibility of the authors and does not necessarily represent the official views of the National Institutes of Health.

## Notes

### Competing Interest Statement

The authors have declared no competing interest.

